# Targeting phosphatase DUSP22 ameliorates skeletal muscle wasting via Akt independent JNK-FOXO3a repression

**DOI:** 10.1101/2024.04.08.588643

**Authors:** Sang-Hoon Lee, Hyun-Jun Kim, Seon-Wook Kim, Hyunju Lee, Da-Woon Jung, Darren Reece Williams

**Affiliations:** New Drug Targets Laboratory, School of Life Sciences, Gwangju Institute of Science and Technology, Gwangju, Jeollanam-do, 61005, Republic of Korea; School of Electrical Engineering and Computer Science, Gwangju Institute of Science and Technology, Gwangju, Jeollanam-do, 61005, Republic of Korea

**Keywords:** Skeletal muscle wasting, DUSP22, sarcopenia, glucocorticoids, immobilization, myofiber atrophy, FOXO3a

## Abstract

Skeletal muscle wasting results from numerous conditions, such as sarcopenia, glucocorticoid therapy or intensive care. It prevents independent living in the elderly, predisposes to secondary diseases, and ultimately reduces lifespan. There is no approved drug therapy and the major causative mechanisms are not fully understood. Dual specificity phosphatase 22 (DUSP22) is a pleiotropic signaling molecule that plays important roles in immunity and cancer. However, the role of DUSP22 in skeletal muscle wasting is unknown. In this study, DUSP22 was found to be upregulated in sarcopenia patients and models of skeletal muscle wasting. DUSP22 knockdown or pharmacological inhibition prevented multiple forms of muscle wasting. Mechanistically, targeting DUSP22 suppressed FOXO3a, a master regulator of skeletal muscle wasting, via downregulation of the stress-activated kinase JNK, which occurred independently of aberrant Akt activation. DUSP22 targeting was also effective in human skeletal muscle cells undergoing atrophy. In conclusion, phosphatase DUSP22 is a novel target for preventing skeletal muscle wasting. The DUSP22-JNK-FOXO3a axis could be exploited to treat sarcopenia or related aging disorders.

## Introduction

Skeletal muscle wasting is a complex multifactorial disorder. A major type of muscle wasting is the loss of mass and strength due to aging, which was termed sarcopenia (“flesh” and “poverty” in Greek) by Irwin Rosenberg in 1989 [1]. It is estimated that skeletal muscle loses around 1% mass and 3% strength annually from middle age, leading to an overall mass reduction that can reach 50% by the 8-9^th^ decade of life [2, 3]. The decreased physical function and increased frailty progressively leads to impaired mobility, loss of independence, and increased mortality. Sarcopenia has become a major health issue due to demographic aging, with an economic burden estimated at $40.4 billion per year in the United States alone [4–7].

In addition to aging, many other disorders can produce skeletal muscle wasting. Examples include side effects from commonly prescribed drugs (e.g. glucocorticoid therapy), infections (such as HIV-AIDS), various degenerative diseases (including heart failure, chronic kidney disease and diabetes), stroke, burns, or intensive care (also known as ICU-acquired weakness) [1]. Currently, there are no clinically approved drugs for treating skeletal muscle wasting and the causative molecular mechanisms are not completely understood [8].

Dual-specificity phosphatases (DUSPs) are a family of signaling enzymes that dephosphorylate serine/threonine and tyrosine residues [9]. Numerous protein and non-protein substrates, such as lipids and glucans, are modified by DUSPs [10, 11]. DUSP22 was recently linked to aging processes in kidney tissue and DUSP22 is also known to be expressed in skeletal muscle [12, 13]. A major target of DUSP22 is the stress-activated c-Jun N-terminal kinase (JNK) signaling cascade that is associated with a numerous diseases, such as cancer, dementia autoimmunity [9, 14–17]. The JNK pathway has also been linked to some types of muscle wasting [18]. JNK signaling is increased in cancer-associated skeletal muscle loss and upregulates the atrogenes atrogin-1 (Fbxo32) MuRF-1 (Trim63) and [18]. JNK signaling has been shown to induce the activation of FOXO3a, which is a master regulator of skeletal muscle wasting and transcriptional activator of MuRF-1 and atrogin-1 [19–22]. Thus, JNK could be a drug target for this disorder. However, despite much research progress, no JNK inhibitor compound has been approved for clinical use [23]. FOXO3a signaling can also be downregulated via activation of the important signaling molecule Akt (protein kinase B), although Akt activating compounds could increase the incidence of aging-related pathologies and may be unsuitable for therapeutic applications [24–26]. Therefore, discovering novel signaling molecules and targets that suppress FOXO3a could facilitate therapeutic development for muscle wasting. However, the potential role of DUSP22-JNK signaling in the pathogenesis of skeletal muscle wasting is unknown.

In this study, DUSP22 expression levels were analyzed in sarcopenia patient samples and experimental models of skeletal muscle wasting. The effect of DUSP22 overexpression, knockdown, and pharmacological targeting with the small molecule inhibitor BML-260 were assessed in cell-based models of muscle atrophy. Skeletal muscle knockdown of DUSP22 and pharmacological inhibition was then investigated in three models of muscle wasting. Mechanism of action was elucidated by gene expression analysis of atrophy-related genes and whole genome transcriptome sequencing. Applicability to humans was validated using donor-derived skeletal muscle cells undergoing atrophy.

## Results

### DUSP22 is upregulated in skeletal muscle wasting and overexpression disrupts myogenesis

The expression of DUSP22 in humans with skeletal muscle wasting was investigated using a sarcopenia gene expression database of human muscle biopsies from elderly individuals over 70 years old and across ethnicities (Singapore Sarcopenia Study; (GEO accession no. GSE111016 [27]). DUSP22 expression was found to be upregulated in sarcopenia patients (Figure 1A). The Dex treatment model of myotube atrophy was used to study the effect of DUSP22 on muscle wasting in vitro, as previously described [28]. DUSP22 expression was upregulated in myotubes undergoing Dex-induced atrophy, as assessed by RNA Seq (Figure 1B) and qPCR (Figure 1C). DUSP22 expression was also investigated in the tibialis anterior (TA) muscle of three models of skeletal muscle wasting: Dex treatment, aged mice (27 months-old), and immobilization (Figure 1C). The mean DUSP22 expression level was found to be upregulated in all three models, and reached statistical significance in the aging and immobilization models.

**Figure 1:**
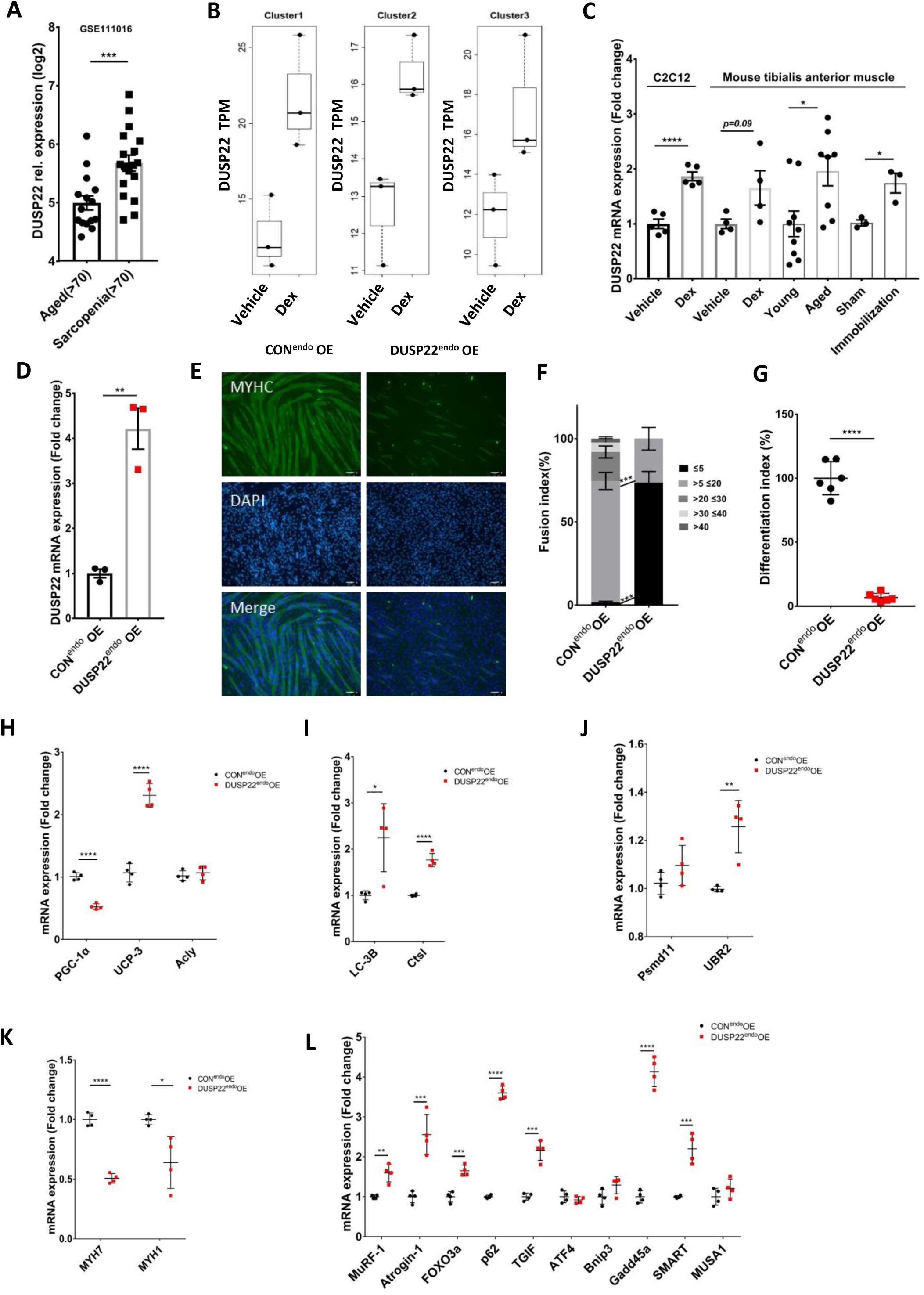
DUSP22 is upregulated in skeletal muscle wasting and overexpression disrupts myogenesis. A) DUSP22 expression in aged individuals (over 70 years) and aged individuals diagnosed with sarcopenia (obtained from the Singapore Sarcopenia Study; (GEO accession no. GSE111016 [27]). B) DUSP22 expression in C2C12 murine myotubes treated with vehicle or dexamethasone (Dex) to induce atrophy. Expression was measured using RNA Seq. TPM=transcript per million. C) qPCR analysis of DUSP22 expression in four models of muscle atrophy: 1) C2C12 myotubes treated with Dex, 2) the TA muscle of C57BL/6 mice treated with Dex, 3) the TA muscle of young (5 months-old) and geriatric (27 months-old) C57BL/6 mice, 4) the TA of C57BL/6 mice after hind limb immobilization. D) qPCR of DUSP22 expression in C2C12 myoblasts transfected with a DUSP22 CRISPR activation plasmid (DUSP22^endo^OE) or control plasmid (CON^endo^OE). E) MYH2 immunocytochemistry of CON^endo^OE and DUSP22^endo^OE myoblasts after 96 h culture in DM (scale bar=100 µm). F) Fusion index. G) Differentiation index. H-L) qPCR analysis of gene expression related to the following: H) Mitochondrial homeostasis (PGC-1α (peroxisome proliferator-activated receptor gamma coactivator 1-alpha), UCP-3 (mitochondrial uncoupling protein 3), Acly (ATP citrate lyase)). I) Autophagy (LC-3B (microtubule-associated proteins 1A/1B light chain 3B), CtsL (cathepsin L)). J) Ubiquitin-proteasome system (UPS) (UBR2 (ubiquitin protein ligase E3), Psmd11 (proteasome 26S subunit, non-ATPase 11). K) Myosin heavy chain levels (MYH7 (β-myosin heavy chain), MYH1 (striated muscle myosin heavy chain 1)), and L) FoxO3a-related signalling (FoxO3a, MurF-1, atrogin-1, p62, TGIF (TGFB induced factor homeobox 1), ATF4 (activating transcription factor 4), Bnip3 (BCL2/adenovirus E1B 19 kDa protein-interacting protein 3); Gadd45a (growth arrest and DNA damage inducible alpha), SMART (specific of muscle atrophy and regulated by transcription), MUSA1 (muscle ubiquitin ligase of SCF complex in atrophy-1)). *=*p*<0.05, **=*p*<0.01, ***=*p*<0.001 and ****=*p*<0.0001 indicate significantly increased or decreased.

DUSP22 was overexpressed in C2C12 myoblasts using the endogenous CRISPR-cas9 editing tool and confirmed by qPCR (Figure 1D). Overexpression disrupted myotube formation and significantly reduced the fusion and differentiation indexes (Figure 1E-F). Expression analysis was carried out for genes linked to mitochondrial function, autophagy, ubiquitin-proteasome system (UPS), myosin heavy chain (MHC) isoforms, and FOXO3a signaling. DUSP22 overexpression down-regulated peroxisome proliferator-activated receptor gamma coactivator 1-alpha (PGC-1α), a master regulator of mitochondrial biogenesis, and upregulated the mitochondrial uncoupling protein UCP-3, although acyl-CoA synthetase long chain family member 1 (Acyl), which regulates mitochondrial fatty acid metabolism, was unaffected (Figure 1H). The autophagy genes microtubule-associated proteins 1A/1B light chain 3B (LC-3B) and cathepsin L1 (Ctsl) were both upregulated by DUSP22 overexpression (Figure 1I). The UPS gene ubiquitin protein ligase E3 component N-recognin 2 (UBR2) was upregulated by DUSP22 overexpression, while proteasome 26S Subunit, Non-ATPase 11 (Psmd11) expression was unaffected (Figure 1J). The MHC isoforms MYH7 and MYH1 were downregulated by DUSP22 overexpression (Figure 1K). 10 genes associated with FOXO3a signaling were analyzed (MuRF-1, atrogin-1, FOXO3a, sequestosome 1 (p62), TGFB induced factor homeobox 1 (TGIF), activating transcription factor 4 (ATF4), BCL2/adenovirus E1B 19 kDa protein-interacting protein 3 (Bnip3), growth arrest and DNA damage inducible alpha (Gadd45a), specific of muscle atrophy and regulated by transcription (SMART), and muscle ubiquitin ligase of SCF complex in atrophy-1 (MUSA1)). Seven genes were found to be upregulated by DUSP22 overexpression (MuRF-1, atrogin-1, FOXO3a, p62, TGIF, Gadd45a, and SMART) (Figure 1L).

### DUSP22 expression correlates with atrogene expression and knockdown prevents myotube atrophy

The potential relationship between DUSP22 and atrogene expression levels were investigated by expression array profiling of a database obtained from muscle biopsies from young and older adults (GSE117525 [29]). The E3 ubiquitin ligase atrogenes MuRF-1 (TRIM63) and atrogin-1 (FBXO32, MAFbx) were selected because they are commonly measured in studies of skeletal muscle wasting and transcriptionally activated by FOXO3a [30]. Expression profiling confirmed a significant positive correlation between the expression of DUSP22 and MuRF-1 or atrogin-1 (Figure 2A-B).

**Figure 2:**
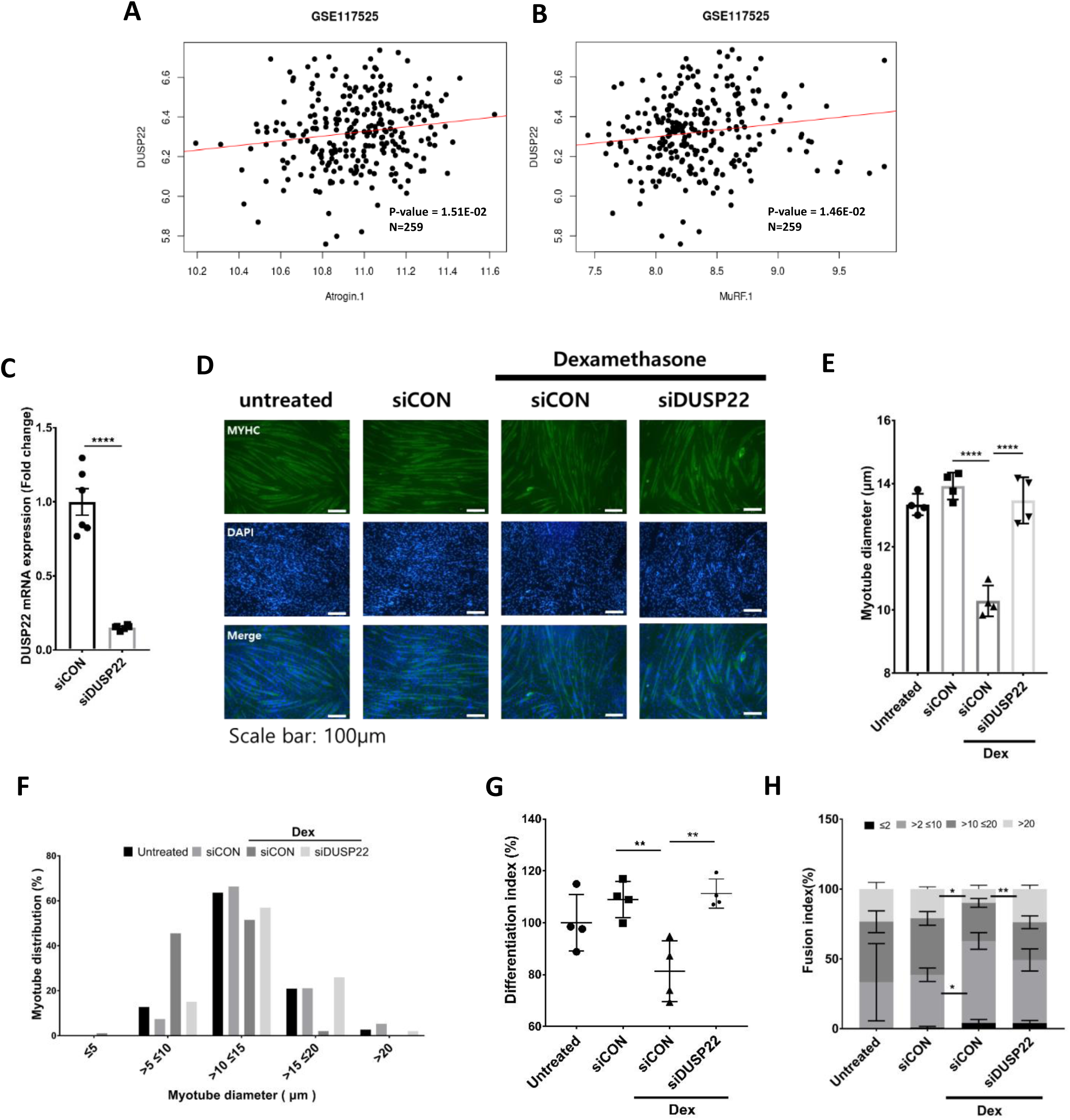
DUSP22 expression positively correlates with MuRF-1 and atrogin-1 expression and knockdown inhibits myotube atrophy. A-B) Expression array profiling of DUSP22 and MuRF-1 and atrogin-1 expression levels in a database obtained from muscle biopsies taken from young and older adults (GSE117525 [37]). C) qPCR analysis of DUSP22 expression in C2C12 myoblasts cultured treated with DM and control siRNA or DUSP22 siRNA for 96 h. D) MYH2 immunocytochemistry of C2C12 myoblasts cultured as follows: (1) 120 h incubation with DM; (2) 72 h incubation with DM and 48 h incubation with DM plus control, DUSP22 siRNA; (3) Following 72 h incubation with DM, 24 h incubation in with DM plus control, scrambled siRNA and additional 24 h treatment with 10 μM Dex plus siRNA; (4) Following 72 h incubation with DM, 24 h incubation in with DM plus DUSP22 siRNA and additional 24 h treatment with 10 μM Dex plus siRNA (scale bar=100 μm). E) Myotube diameter. F) Myotube distribution. (n=6). G) Differentiation index of C2C12 myoblasts treated as in part (C). H) Fusion index. *=*p*<0.05, **=*p*<0.01, ***=*p*<0.001 and ****=*p*<0.0001 indicate significantly increased or decreased.

siRNA was used to investigate the effect of DUSP22 knockdown on myotube atrophy and atrogene expression. Gene knockdown in the myotubes was confirmed by qPCR (Figure 2C). Knockdown prevented atrophy in the Dex model, as shown by increased myotube diameter and a greater proportion of larger-sized myotubes (Figure 2D-F). Knockdown also increased the myotube fusion and differentiation indexes (Figure 2G-H).

FOXO3a signalling activity is inhibited via phosphorylation by activated Akt (also known as protein kinase B), which stimulates muscle hypertrophy [30]. FOXO3a phosphorylation in myotubes was decreased by Dex and increased by DUSP22 knockdown (Figure 2G-H).

### DUSP22 knockdown lowers atrogene expression, downregulates JNK, and enhances myogenesis

The activation of FOXO3a and Akt after DUSP22 knockdown was assessed by western blotting. Knockdown in normal and Dex treated myotubes increased FOXO3a phosphorylation, indicating reduced signaling activity (Figure 3A-B). Knockdown had no significant effect on Akt phosphorylation (Figure 3A-C). qPCR and western blotting showed that DUSP22 knockdown reduced atrogin-1 and MuRF-1 levels in Dex treated myotubes (Figure 3D-F).

**Figure 3:**
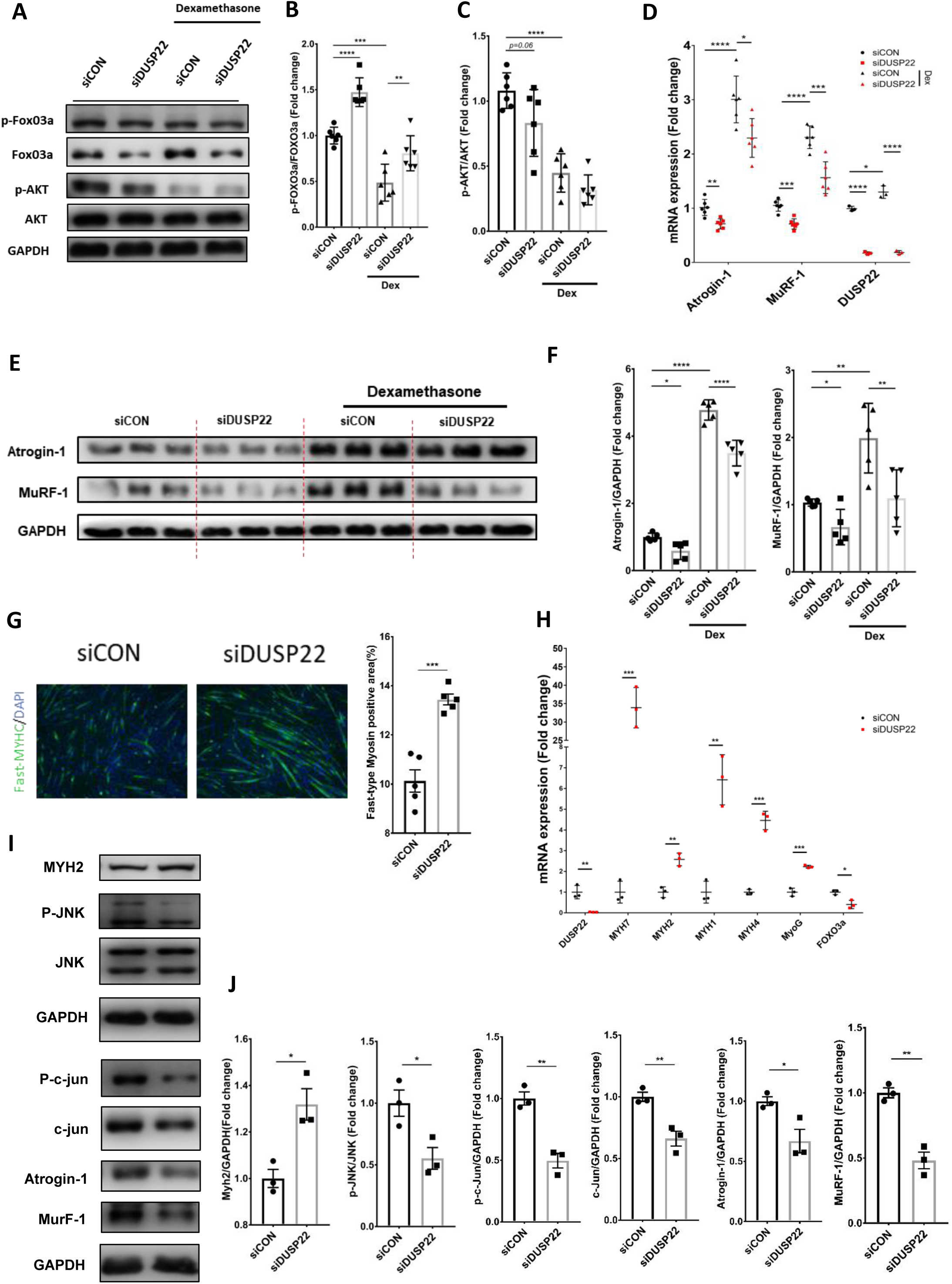
DUSP22 knockdown inhibits atrogene expression and promotes myogenesis. A) Western blot analysis of FoxO3a and Akt phosphorylation in C2C12 myotubes treated with control or DUSP22 siRNA in the presence or absence of Dex. B) Quantification of FOXO3a phosphorylation, which is inversely proportional to activity. C) Quantification of Akt phosphorylation, which is directly proportional to activity. D) qPCR analysis of atrogin-1, MuRF-1, and DUSP22 expression. E) Western blot of atrogin-1 and MuRF-1 expression levels. F) Quantification of atrogin-1 and MuRF-1 levels. G) Fast-type myosin immunostaining of C2C12 myoblasts after culture in DM for 24 h and treatment with control, scrambled siRNA or DUSP22 siRNA for 72 h. Quantification of the fast-type myosin positive myotubes is also shown. H) qPCR analysis of DUSP22, myosin heavy chains (MYH7, MYH2, MYH1, and MYH4), myogenin (MyoG) and FOXO3a expression. I) Western blot analysis of MYH2, JNK, c-jun, c-jun phosphorylation, atrogin-1 and MuRF-1. J) Quantification of expression and phosphorylation levels. *=*p*<0.05, **=*p*<0.01, ***=*p*<0.001 and ****=*p*<0.0001 indicate significantly increased or decreased.

The effects of DUSP22 knockdown was further investigated in differentiating myoblasts under normal conditions. Knockdown increased the expression of fast type myosin in the myotubes (Figure 3G). qPCR analysis showed that knockdown increased expression of the myosin heavy chains MYH7, MYH2, MYH1, and MYH4 (Figure 3H). In addition, DUSP22 knockdown increased the expression of myogenin (MyoG), a master inducer of myogenesis, and decreased the expression of FOXO3a (Figure 3H). Western blotting indicated that DUSP22 knockdown increased MYH2 mean expression, although it did not reach statistical significance (Figure 3I-J). Western blotting also confirmed that DUSP22 downregulates the JNK pathway in muscle cells. Knockdown decreased JNK levels and the phosphorylation and expression of the downstream JNK pathway mediator, c-jun (Figure 3I-J). As previously observed in Figure 3E, DUSP22 knockdown also reduced atrogin-1 and MuRF-1 expression levels in normal myotubes (Figure 3I-J).

### DUSP22 pharmacological targeting prevents myotube atrophy

BML-260 is a rhodanine-based small molecule inhibitor of DUSP22 [31]. Molecular docking analysis indicated that BML-260 non-covalently binds to the active site of human DUSP22 at residue Cys88 with a Vina score of -5.8 (Figure 4A). C2C12 mouse myotubes were treated with Dex in the presence or absence of BML-260. Dex treated myotubes underwent atrophy, as shown by a reduction in myotube diameter. BML-260 treatment prevented myotube atrophy and maintained the proportion of larger-sized myotubes (Figure 4B-D). BML-260 also recovered the myotube fusion and differentiation indexes (Figure 4E-F). In addition, BML-260 prevented the reduction in protein synthesis caused by Dex (Figure 4G-H). qPCR analysis showed that atrogin-1 and MuRF-1 were upregulated by Dex treatment and downregulated by BML-260 (Figure 4I). Western blotting confirmed that BML-260 reduced atrogin-1 and MuRF-1 levels in the Dex treated myotubes (Figure 4J-L).

**Figure 4:**
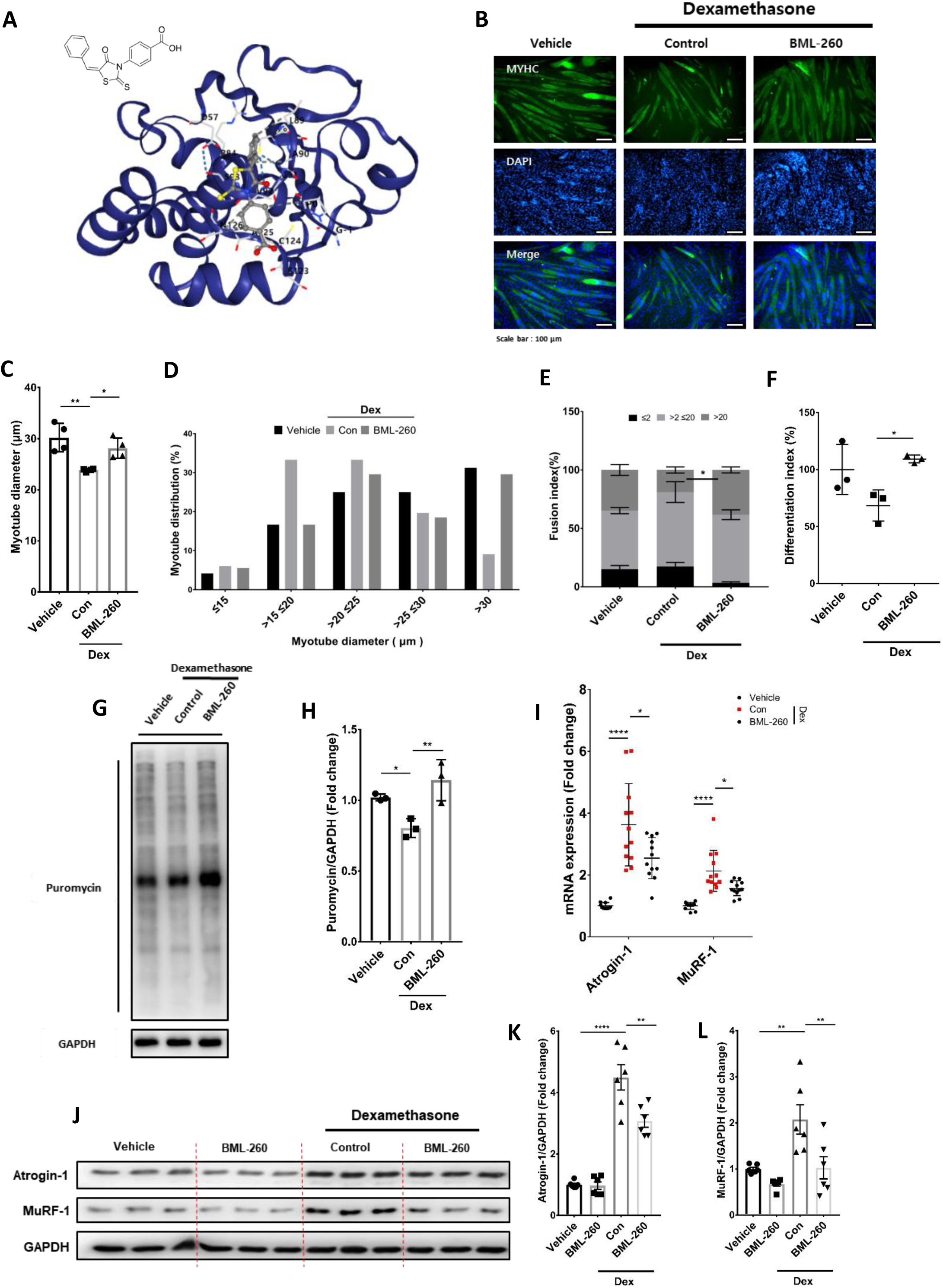
DUSP22 pharmacological targeting prevents myotube atrophy. A) Chemical structure of the DUSP22 inhibitor, BML-260, and CB-Dock2 modeling of BML-260 binding to the active site of human DUSP22 (Pocket C5 and score -5.8. Chain A: GLY-1, PRO0, ASP57, CYS88, LEU89, ALA90, GLY91, VAL92, SER93, ARG94, SER123, CYS124, ALA125, ASN126, ASN128). B) MYH2 immunostaning of C2C12 myoblasts cultured as follows: (1) DM for 120 h (untreated); (2) DM for 96 h and DM plus 10 μM Dex for 24 h; (3) DM for 96 h and DM plus 10 μM Dex and 12.5 μM BML-260 for 24 h (scale bar=100 μm). C) Mean myotube diameter. D) Myotube diameter distribution. E) Fusion index. F) Differentiation index. G) SUnSET assay of protein synthesis rate measuring puromycin incorporation. H) Quantification of the SUnSET assay. I) qPCR analysis of atrogin-1 and MuRF-1 expression. J) Western blot analysis of atrogin-1 and MuRF-1. K-L) Quantification of atrogin-1 and MuRF-1 levels. *=*p*<0.05, **=*p*<0.01, and ****=*p*<0.0001 indicate significantly increased or decreased.

### DUSP22 knockdown in skeletal muscle ameliorates wasting

DUSP22 siRNA was delivered to the TA muscle of Dex treated mice (Figure 5A). DUSP22 knockdown in the TA was confirmed by western blotting (Figure 5B-C). DUSP22 knockdown prevented TA muscle wasting (Figure 5D). Histological analysis indicated that knockdown increased myofiber CSA and the proportion of larger sized myofibers (Figure 5E-G). Fast twitch type 2 myofibers are known to be more susceptible to sarcopenia [32]. DUSP22 knockdown preserved the CSA of type 2a and type 2b myofibers (Figure 5E, H-I). Western blotting showed that FOXO3a activity increased after Dex treatment and was reduced by DUSP22 knockdown (Figure 5J-K). DUSP22 knockdown also downregulated JNK and the downstream effector, c-jun (Figure 5J-L). The upregulation of atrogin-1 and MuRF-1 by Dex was also inhibited by DUSP22 knockdown (Figure 5M-N). A similar result was observed for p62, a key autophagy-related gene induced by FOXO3a [33] (Figure 5M-N).

**Figure 5:**
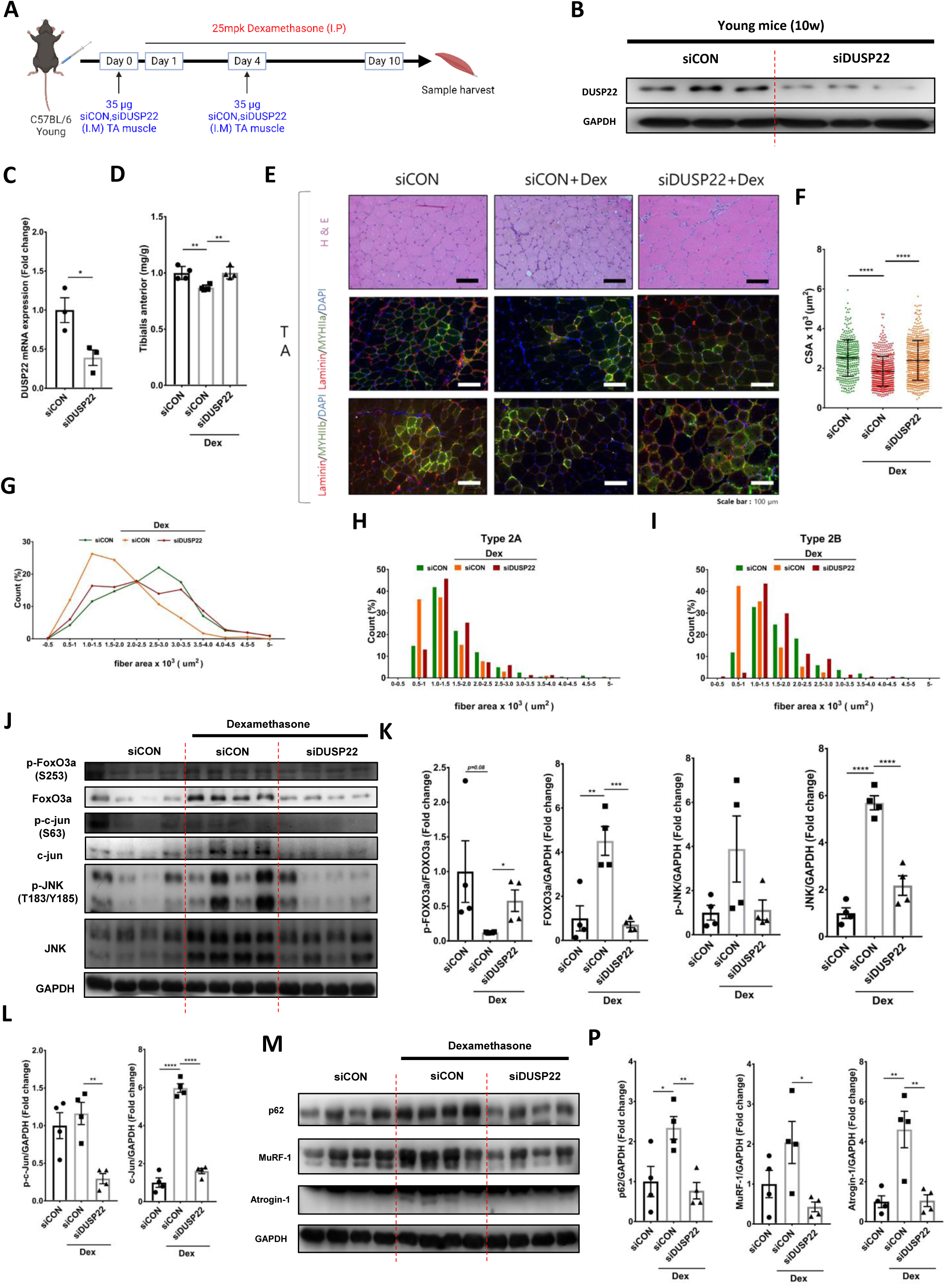
DUSP22 knockdown prevents Dex-induced skeletal muscle atrophy. A) Schematic of the experimental protocol. B) Western blot analysis of DUSP22 in the Dex-treated tibialis anterior (TA) muscle 3d after delivery of control or DUSP22 siRNA. C) Quantification of DUSP22 expression. D) TA muscle mass. E) Representative H&E staining, and myosin heavy chain IIa (MYHIIa; type 2A) and IIb (MYHIIb; type 2B) immunostaining of the TA muscle. F) TA myofiber CSA. G) TA myofiber area distribution. H) Type 2A myofiber distribution. I) Type 2B myofiber distribution. J) Western blot analysis of phosphorylated FOXO3a (P-FOXO3a), FOXO3a, phosphorylated c-jun (P-c-jun), c-jun, phosphorylated JNK (P-JNK), JNK in the TA muscle. GAPDH was used for normalization of expression. K) Quantification of P-FOXO3a, FOXO3a, P-JNK, and JNK. L) Quantification of P-c-jun and c-jun. M) Western blot analysis of p62, MuRF-1, and atrogin-1. GAPDH was used for normalization of expression. N) Quantification of p62, MuRF-1, and atrogin-1. *=*p*<0.05, **=*p*<0.01, ***=*p*<0.001 and ****=*p*<0.0001 indicate significantly increased or decreased.

### DUSP22 pharmacological targeting ameliorates skeletal muscle wasting

The Dex treatment mouse model was used to investigate DUSP22 pharmacological targeting with BML-260 (Figure 6A). Dex significantly reduced mean body weight, which was recovered by BML-260 treatment (Figure 6B). BML-260 treatment increased the mean grip strength value, but it did not achieve statistical significance, possibly due to variation in the Dex alone group (Figure 6C). Rotarod testing showed that BML-260 significantly recovered muscle performance in both the acceleration and constant models (Figure 6D). Histological analysis of the TA muscle showed that BML-260 significantly increased myofiber CSA (Figure 6E-F). BML-260 also shifted the myofiber area distribution towards larger sized fibers (Figures 6G). TA muscle mass was increased by BML-260 treatment (Figure 6H), and the upregulation of atrogin-1 and MuRF-1 was inhibited (Figure 6I-J).

**Figure 6:**
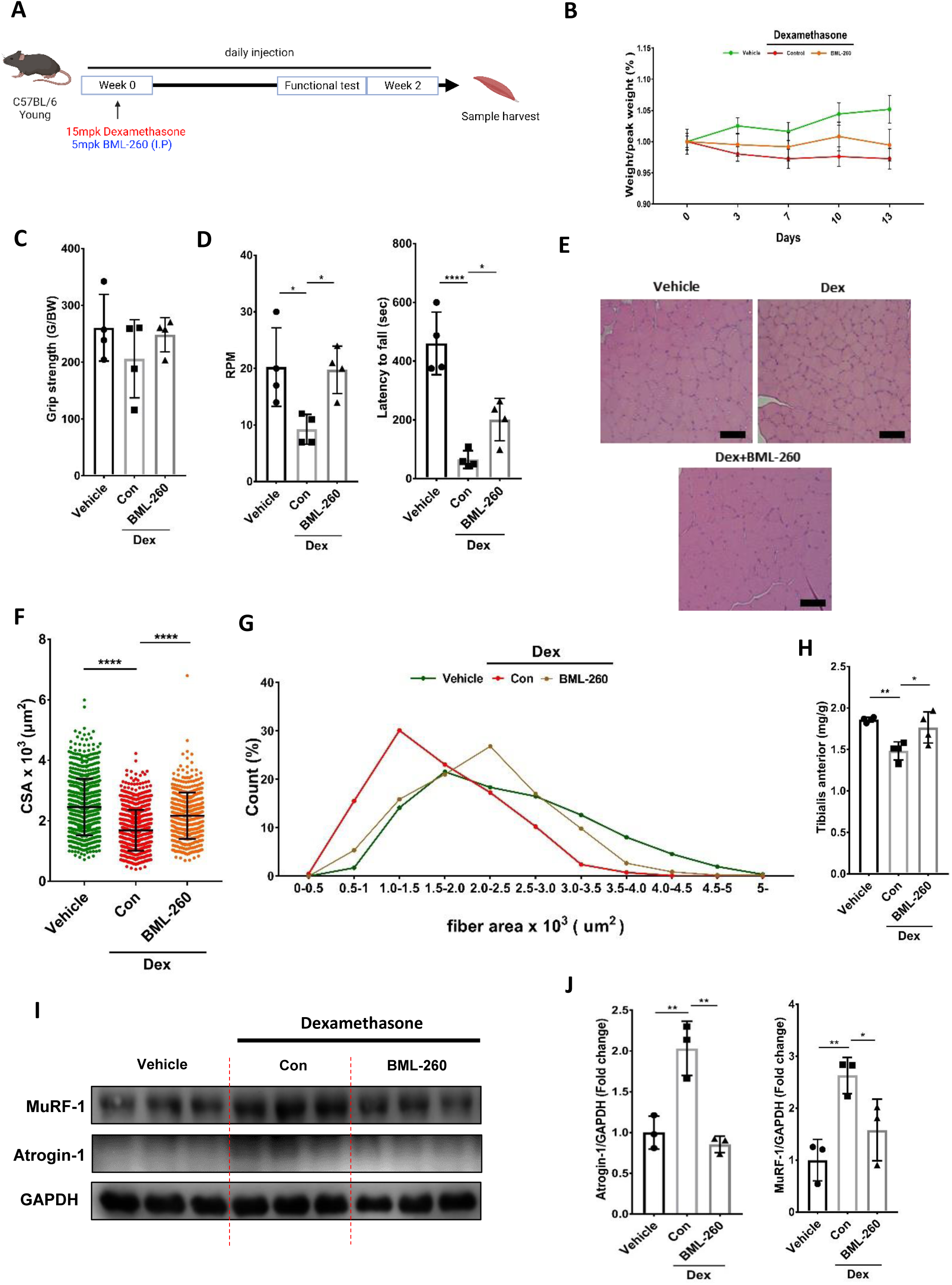
DUSP22 pharmacological targeting prevents Dex-induced skeletal muscle atrophy. A) Schematic of the experimental protocol. B) Body weight during 13 d treatment with vehicle, 15 mg/kg Dex, or 15 mg/kg Dex and 5 mg/kg BML-260. C) Grip strength. D) Rotarod performance in the constant (RPM) and acceleration (latency to fall) models. E) Representative H&E staining of the gastrocnemius muscle. F) Myofiber CSA. G) Myofiber area distribution. H) TA muscle mass. I) Western blot analysis of atrogin-1 and MuRF-1 expression in the gastrocnemius muscle. GAPDH was used for normalization of expression. *=*p*<0.05, **=*p*<0.01, and ****=*p*<0.0001 indicate significantly increased or decreased.

### DUSP22 knockdown and pharmacological inhibition prevents aging-induced skeletal muscle wasting

DUSP22 siRNA was delivered to the TA muscle of geriatric mice (27 months-old) (Figure 7A). DUSP22 knockdown in the TA was confirmed by western blotting (Figure 7B-C). DUSP22 knockdown also reduced the upregulation of MuRF-1 (Figure 7B). TA muscle mass was increased in the DUSP22 knockdown group (Figure 7D). Histological analysis revealed that DUSP22 knockdown increased TA myofiber CSA (Figure 7E-F). Western blot analysis showed that atrogin-1, MuRF-1, and FOXO3a levels tended to be reduced in the siRNA treated TA muscle compared to the contralateral leg (7G-H).

**Figure 7:**
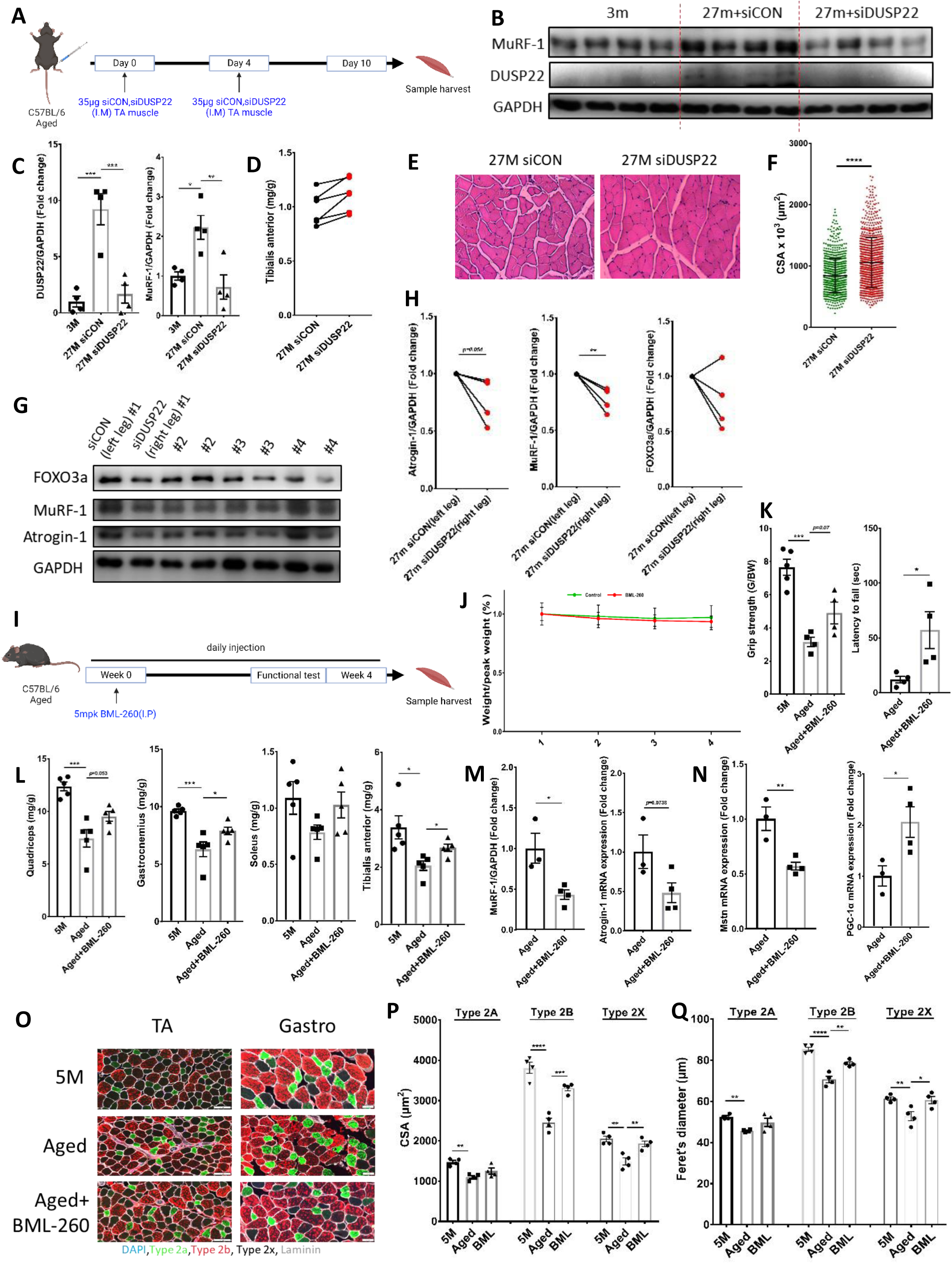
DUSP22 knockdown or pharmacological protects against aging-induced skeletal muscle wasting. A) Experimental protocol to knockdown DUSP22 expression in the TA of aged mice. B) Western blot analysis of DUSP22 and MuRF-1 and expression levels. GAPDH was used for normalization of expression. C) Quantification of DUSP22 and MuRF-1. D) Change in TA muscle mass over the course of the experiment. E) Representative H&E staining of the TA muscle. F) TA myofiber CSA. G) Western blot of FOXO3a, atrogin-1, and MuRF-1 levels in the TA muscle from the control siRNA treated left and DUSP22 siRNA treated right leg. H) Change in FOXO3a, atrogin-1, and MuRF-1 expression relative to GAPDH. I) Experimental protocol for DUSP22 pharmacological targeting in aged mice. J) Body weight over the course of the experiment. K) Grip strength and rotarod performance in the latency to fall test. L) Mass of the quadriceps, gastrocnemius, TA and soleus muscles. M) qPCR analysis of MuRF-1 and atrogin-1 expression. N) qPCR analysis of myostatin (Mstn) and PGC-1α expression. O) MYHIIa (type 2a), MYHIIb (type 2b), and MYHIIx (type 2x), DAPI and laminin staining in the TA and gastrocnemius muscles (5M=5 months-old). Scale bar=100 µm. P) CSA of the type 2a, 2b, and 2x myofibers. O) Minimal Feret’s diameter of the type 2a, 2b, and 2x myofibers. *=*p*<0.05, **=*p*<0.01, ***=*p*<0.001 and ****=*p*<0.0001 indicate significantly increased or decreased.

Geriatric mice (24-26 months-old) were treated with BML-260 for 4 weeks (Figure 7I). Body weight was not significantly affected by BML-260 treatment (Figure 7J). The grip strength mean value was increased by BML-260 treatment but did not reach statistical significance (Figure 7K). Latency to fall in the rotarod test was statistically increased by BML-260 treatment (Figure 7K). BML-260 also increased the mean mass value of the quadriceps, gastrocnemius, TA and soleus muscles, and reached statistical significance for the TA and gastrocnemius muscles (Figure 7L). qPCR analysis indicated that BML-260 reduced the expression of atrogin-1 and MuRF-1 in the TA muscle (Figure 7M). BML-260 also reduced the expression of myostatin and increased the expression of PGC-1α, which are critical promoters and inhibitors of muscle aging, respectively (Figure 7N). Immunostaining of the TA and gastrocnemius muscles showed that BML-260 increased the CSA and Feret’s diameter of the fast twitch type 2A, type 2B and 2X myofibers in the aged mice (Figure 7O-Q).

### DUSP22 pharmacological targeting in aged skeletal muscle regulates FOXO3a signaling, muscle cell differentiation and myofiber development

The mechanisms by which DUSP22 targeting prevents aging-induced skeletal muscle atrophy were characterised by whole genome RNA seq of the TA muscle in geriatric mice (27 months-old). Principal component analysis showed that pharmacological targeting with BML-260 produced similar overall effects on gene expression in the aged mice (Figure 8A). The volcano plot showed a total of 350 differentially expressed genes (DEGs) (Figure 8B). DUSP22 pharmacological targeting downregulated genes linked to FOXO3a signalling (including atrogin-1 and MuRF-1), as assessed by overall TPM fold change (Figure 8C) and normalized counts from the raw read (Figure 8D). Interestingly, pharmacological targeting also upregulated the expression of musclin (OSTN), which is an exercise-induced myokine that protects against muscle wasting [34, 35] (Figure 8E). Gene ontology (GO) analysis showed that pharmacological targeting upregulated genes linked to muscle cell differentiation and muscle fiber development (Figure 8F). In addition to musclin, pharmacological targeting increased the expression of Myh4 (myosin heavy chain 2b), which is a predominant myosin type in fast-twitch TA muscle (Figure 8F). The genes Six1 and Six4, which are known to activate the fast-type muscle gene program [36], were also upregulated (Figure 8F). Genes linked to the PI3K-Akt pathway were also downregulated by pharmacological targeting (Figure 8G). Gene set enrichment analysis (GSEA) combined with GO analysis indicated that skeletal muscle genes involved in the negative regulation of hypertrophy and response to inactivity were under-represented after DUSP22 targeting (Figure 8H). In addition, genes involved in skeletal muscle cell differentiation and proliferation were enriched after targeting (Figure 8I).

**Figure 8:**
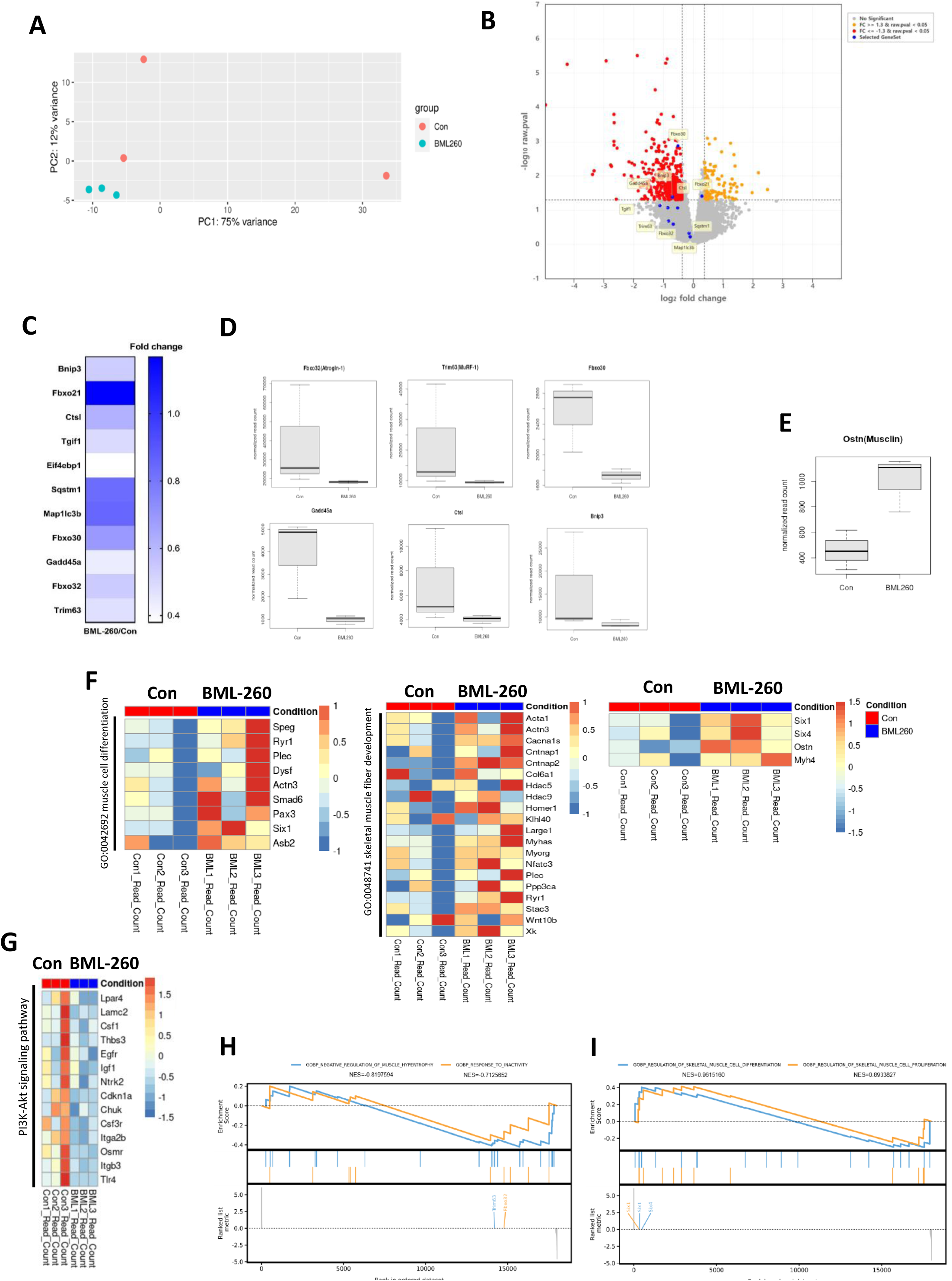
Whole genome RNA Seq analysis of DUSP22 pharmacological targeting in geriatric mouse TA muscle. A) Principal component analysis of gene expression variance in the vehicle treated (Con) and BML-260 treated groups. B) Volcano plot of differentially expressed genes. Selected genes linked to FOXO3a signaling are indicated. C) Expression analysis of genes related to FOXO3a signaling measured by overall TPM fold change. D) Expression of selected genes linked to FOXO3a signaling as normalized counts from the raw read. E) Expression of the exercise-induced myokine musclin (Ostn). F) Gene ontology (GO) analysis of the cellular components muscle cell differentiation and muscle fiber development. Also shown are Six1 and Six4 (which specify fast twitch myofiber development), Ostn and Myh4 (myosin heavy chain 2b; the predominant myosin type in the fast-twitch TA muscle). G) Expression changes for genes linked to PI3K-Akt signaling. H) Gene set enrichment analysis (GSEA) combined with GO analysis of genes involved in the negative regulation of muscle hypertrophy and response to inactivity. The rankings for atrogin-1 (Fbox32) and MuRF-1 (Trim63) are indicated. I) (GSEA) combined with GO analysis of genes involved in the regulation of skeletal muscle differentiation and proliferation. The rankings for Six1 and Six4 are indicated.

### DUSP22 pharmacological targeting downregulates atrogenes in immobilized muscle and ameliorates atrophy in human skeletal myotubes

A hind limb immobilization model was used to assess whether DUSP22 pharmacological targeting is effective for multiple types of skeletal muscle wasting. Immobilization decreased the mean mass of the quadriceps, gastrocnemius, TA and soleus muscles, and reached statistical significance for the gastrocnemius (Figure 9A). Analysis of the TA muscle showed an upregulation of DUSP22 expression, which was inversely correlated with TA mass (Figure 9B-C). BML-260 treatment increased the mean TA mass, although it did not reach statistical significance (Figure 9D). Atrogin-1 and MuRF-1 expression was upregulated in the immobilized TA and downregulated by BML-260 treatment (Figure 9E).

**Figure 9:**
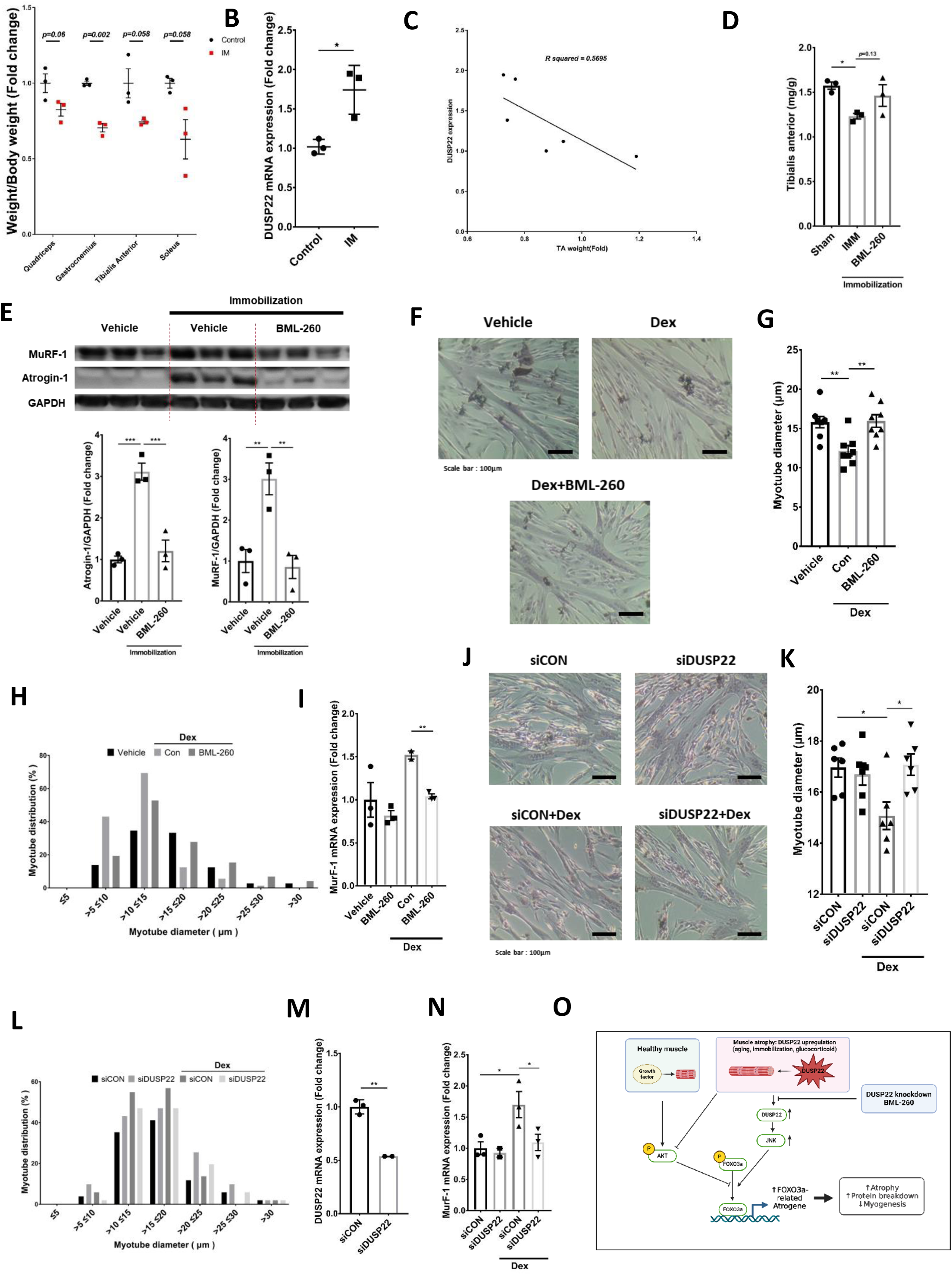
DUSP22 targeting downregulates atrogene expression in immobilized skeletal muscle and inhibits atrophy in human donor myotubes. A) Effect of immobilization on skeletal muscle mass using the plastic EP tube method. B) qPCR analysis of DUSP22 expression in the TA muscle (IM=immobilized). C) Plot of DUSP22 expression in relation to TA mass. D) TA mass. E) Western blot analysis of atrogin-1 and MuRF-1 levels in the TA muscle. Quantification of atrogin-1 and MuRF-1 levels relative to GAPDH are also shown. F) Micrographs of H&E-stained human myotubes treated as follows: 1) Vehicle alone, 2) 10 μM Dex for 24 h, 3) 10 μM Dex and 12.5 μM BML-260 for 24 h. G) Mean myotube diameter. H) Myotube diameter distribution. I) qPCR analysis of MuRF-1 expression. J) Micrographs of H&E-stained human myotubes treated as follows: (1) 48 h incubation in with DM plus control, scrambled siRNA; (2) 48 h incubation with DM plus DUSP22 siRNA: (3) 24 h incubation with DM plus control, scrambled siRNA and additional 24 h treatment with 10 μM Dex plus siRNA; (4) 24 h incubation with DM plus DUSP22 siRNA and additional 24 h treatment with 10 μM Dex plus siRNA. K) Myotube diameter. L) Myotube diameter distribution. M) qPCR analysis of DUSP22 expression. N) qPCR analysis of MuRF-1 expression. *=*p*<0.05, **=*p*<0.01, and ***=*p*<0.001 indicate significantly increased or decreased. O) Working model of the effect of DUSP22 targeting on skeletal muscle atrophy: 1) In healthy muscle, Akt signaling can promote hypertrophy by increasing protein synthesis and inhibiting the activity of FOXO3a. 2) In the context of skeletal muscle wasting, the Akt pathway can become suppressed and FOXO3a signaling is upregulated. The results from this study show that targeting DUSP22 in wasting muscle downregulates JNK and reduces FOXO3a signaling. These events occur independently of Akt signaling, which remains suppressed. DUSP22 targeting, via pharmacology or gene knockdown, is sufficient to enhance function, improve histopathology, and lower atrogene expression in multiple forms of skeletal muscle wasting.

The clinical potential of DUSP22 targeting for skeletal muscle wasting was tested in cultures of human donor myotubes undergoing Dex-induced atrophy. BML-260 treatment recovered the mean myotube diameter and proportion of larger myotubes (Figure 9F-H). qPCR analysis showed that BML-260 inhibited MuRF-1 expression (Figure 9I), although atrogin-1 expression was not affected (data not shown). DUSP22 gene knockdown also prevented atrophy in human myotubes, as shown by a significant increase in myotube diameter and the proportion of larger sized myotubes (Figure 8J-L). qPCR confirmed the knockdown of DUSP22 (Figure 8M) and showed an inhibitory effect on MuRF-1 upregulation (Figure 8N).

## Discussion

Skeletal muscle wasting has become major socioeconomic problem and there is currently no clinically approved drug for this disorder. In the current study, pharmacological and genetic targeting of the pleiotropic signalling phosphatase DUSP22 was shown to be effective at ameliorating skeletal muscle wasting in multiple models via the suppression of JNK and FOXO3a signalling.

DUSPs are a diverse family of enzymes with at least 61 members in humans [37]. DUSPs have been shown to have key roles in cell growth, differentiation, immunity and neurobiology [38]. DUSP signaling has also been linked to numerous diseases [38]. However, our search of the literature indicates that there are few published studies of the role of DUSP enzymes in skeletal muscle. Haddock, et al., showed that DUSP4 expression was increased in denervation-induced muscle atrophy and, in myotube cultures, reduced extracellular signal-regulated kinase activity [39]. Cooper, et al., reported that DUSP29 expression was also increased in denervation-induced muscle atrophy and lowered glucocorticoid receptor activity in myotube cultures [40]. Pourteymour, et al., showed that the expression patterns of DUSP5 and DUSP6 are oppositely regulated by acute exercise [41]. The finding that DUSP22 targeting is effective at preventing skeletal muscle wasting may provide research impetus to further characterize the potential roles of DUSP family members in the pathogenesis of this disorder.

DUSP22 pharmacological targeting was carried out with the small molecule inhibitor, BML-260. This compound is based on the rhodanine (2-thioxothiazolidin-4-one) chemical scaffold, which is becoming more prominent in medicine and pharmacy [42]. Rhodanine is a privileged heterocyclic compound that is amenable for structural modification and lead development, and has been classified as a highly decorated scaffold with an important position for drug development [43]. Consequently, BML-260 should be suitable for further rounds of structural optimization to improve its drug-like properties, biological stability and targeting of DUSP22. Due to its yellow color and the ability of rhodanine to interact photometrically in biologic assessments [44], it may also be interesting to assess BML-260 as a theranostic agent acting as both a diagnostic biomarker and therapeutic drug for skeletal muscle wasting.

Pharmacological targeting or gene knockdown of DUSP22 inhibited signaling by JNK (also known as stress-activated kinase) and FOXO3a (a key regulator of skeletal muscle atrophy). DUSP22 is also known as JNK-stimulating phosphatase 1 (JSP1) and JNK activation has been shown to promote FOXO3a activation in numerous cell types [19–22]. To confirm the ability of DUSP22 to stimulate JNK in this study, we showed that gene knockdown in skeletal muscle downregulated JNK and the downstream mediator, c-jun. The IGF-1/PI3K/Akt pathway is also known to suppress FOXO3a activity in skeletal muscle and have a major role in the prevention of wasting [45]. However, we consistently found that Akt activity was not upregulated by DUSP22 knockdown or BML-260 treatment. This may be advantageous for developing therapeutic approaches that target FOXO3a independently of the IGF-1-PI3K-Akt pathway due to potential side effects associated with PI3k-Akt activation [46, 47]. In addition, over-activation of protein synthesis, which is a downstream target of IGF-1-PI3K-Akt, has recently been observed in aging-related skeletal muscle wasting [48, 49]. Strategies to further increase protein synthesis in geriatric muscle may worsen proteostatic stress [48]. Compounds such as BML-260, which inhibit FOXO3a signalling via alternative pathways, such as JNK, may be an attractive option for future drug development. A schematic of the effect of DUSP22 targeting in skeletal muscle wasting shown in Figure 8O.

A number of small molecule JNK inhibitors have been developed and are under investigation for disorders such as stroke, cancer and Parkinson’s disease [50, 51]. However, although there has been much research progress, no JNK targeting drug is approved for clinical use. DUSP22 is a relatively unexplored drug target and further investigation may produce lead compounds that can enter clinical trials as indirect JNK inhibitors for a number of disease applications. The in vivo RNA Seq data indicates that Six1 and Six4, genes linked to fast twitch myofiber formation [36], are upregulated by DUSP22 pharmacological targeting in aged muscle. This could explain the increased proportion of fast twitch type 2B and 2X myofibers. The mechanism by which DUSP22 targeting increases fast myofiber number in aged mice (preservation of pre-existing myofibers versus upregulation of transcriptional activators of fast fiber type gene expression) can be an interesting subject for future research. The discovery that musclin is upregulated after DUSP22 targeting may also warrant further investigation, because this myokine has been shown to be critical for cardiac conditioning [52]. Cardiac hypertrophy and skeletal muscle wasting are known to share similar molecular mechanisms, and cardio-sarcopenia has recently been described as a syndrome of concern in aging [53, 54]. In addition, skeletal muscle-secreted musculin has recently been shown to alleviate depression in animal models [55, 56]. To our knowledge, there is no previous report of DUSP22 targeting in cardiac disease or depressive disorders. Whether musclin levels are directly regulated by DUSP22, or via downstream effects on JNK-FOXO3a inhibition, may also be an interesting topic for myokine research.

DUSP22 targeting downregulated the atrogenes atrogin-1 and MuRF-1 in multiple animal models of muscle wasting. However, only MuRF-1 was consistently downregulated in human skeletal myotubes undergoing atrophy. Atrogin-1 and MuRF-1 are E3 ligases and it has previously been shown that MuRF-1 is particularly important in muscle mass homeostasis [57, 58]. MuRF-1 knockout mice are especially resistant to numerous forms of skeletal muscle wasting and it is the only E3 ligase that degrades the α-actin, myosin, and troponin contractile proteins that constitute up to 70% of myofiber protein content [59–61]. Based on these previous reports, it can be envisioned that MuRF-1 downregulation in human muscle cells treated with DUSP22 inhibitors may be sufficient to prevent wasting.

Although atrogin-1 and MuRF-1 are the most commonly assessed E3 ligases in studies of skeletal muscle wasting, other E3 ligases are expressed in skeletal muscle and some of these have been linked to wasting mechanisms. For example, MUSA1 and SMART were shown to participate in skeletal muscle proteolysis [62]. The qPCR analysis presented herein indicates that SMART expression in myotubes increases after DUSP22 overexpression, whereas MUSA1 expression was unchanged. It should be interesting to assess the expression patterns of these additional E3 ligases in more detail after DUSP22 targeting in muscle wasting models.

In summary, this study demonstrates that DUSP22 is a regulator of skeletal muscle wasting and therapeutic target that functions by inhibiting a DUSP22-JNK-FOXO3a signalling axis. These results increase our understanding of the molecular mechanisms driving skeletal muscle wasting and provide a novel candidate for drug development. Previous research has linked aberrant DUSP22 to other diseases, such as T-cell lymphoma and Alzheimer’s disease [63]. The results presented herein may also be applicable to therapeutic approaches for these diseases. In addition, the identification of DUPS22 as a mediator of aging-related skeletal muscle wasting suggests that it may be worthwhile to investigate its role in other aging-related disorders, such as osteoarthritis, heart failure and type 2 diabetes. Overall, these findings provide new insight into the functions of DUSP22, the mechanisms responsible for producing skeletal muscle wasting, and novel treatment approaches for this debilitating condition.

## Materials and methods

### Molecular docking analysis

The binding of BML-260 to human DUSP22 was analysed with the CB-Dock2 software [64]. The Vina score was calculated using DUSP22 human (PDB: 6lvq) and BML-260 (CID: 1565747).

### Cell culture

C2C12 murine skeletal muscle myoblasts cell were purchased from Koram Biotech. Corp, Republic of Korea, and cultured in growth media (GM), containing Dulbecco’s Modified Eagle’s Medium (DMEM), 10% fetal bovine serum, and 1% penicillin and streptomycin (PenStrep). C2C12 myoblasts were differentiated into myotubes at more than 80% confluence by culture in differentiation media (DM), containing DMEM supplemented with 2% horse serum and PenStrep, for 96 h. To induce atrophy, the myotubes were treated with DM plus 10 μM Dex for 24 h.

Human skeletal myoblasts were purchased from Thermo-Fisher Scientific, USA, re-suspended in DM and seeded onto 12-well culture plates at a density of 4.8 × 10^4^ cells/well. 72 h later, the differentiated myotubes were treated with 10 μM Dex for 24 h, stained with haematoxylin and eosin (H&E) using a kit (Merck, Germany) and myotube diameter measured by light microscopy analysis of DIC captured images (Olympus CKX41, Olympus Life Science, Japan) and the NIH imaging software Image J 64 (National Institutes of Health, MD, USA). At least 80 myotubes in 5 micrographs were measured in each experimental group.

### Myotube diameter

Myosin heavy chain 2 (MYH2) immunocytochemistry was used to visualize murine myotubes and measure myotube diameter, following the previously published protocol [65]. After mounting, fluorescence images were captured in 5 different regions using a DMI 3000 B microscope (Leica, Germany), and analysed using the NIH imaging software Image J 64. 100 myotubes containing at least three nuclei were measured in each experimental group.

### Myotube fusion and differentiation indexes

To assess myoblast fusion into multinucleate myotubes, nuclei number within MYH2-positive cells were measured using the NIH imaging software Image J 64. The differentiation index was calculated as the percentage of nuclei in MYH2-positive cells (nuclei in myotubes/total nuclei). Myotubes were classified as MYH-positive cells containing at least 4 nuclei. Myotubes were measured in groups of at least 100, with at least 7500 nuclei measured in each group.

### SUnSET assay

The surface sensing of translation (SUnSET) assay was used to measured protein synthesis, in accordance with the published protocol [66]. In brief, 1 μg/mL puromycin was added to the myotube cultures and cell lysates were then harvested 10 min later. Western blotting was carried out using an anti-puromycin 12D10 antibody (MABE343; Millipore).

### Western blotting

Western blotting was carried out using the previously published protocol [67]. The NIH imaging software Image J 64 was used for densitometry analysis of the gel bands. The primary and secondary antibodies used in this study are shown in reagent and tools table.

### Real-time qPCR

The StepOnePlus Real Time PCR System (Applied Biosystems, UK) was used to measure the mRNA level of the genes of interest. An AccuPower RT PreMix (Bioneer, Republic of Korea) was used to synthesize the cDNA from total RNA. The real-time PCR (qPCR) was carried out according to the manufacturer’s instructions, and the previously published protocol [67]. The primers used in this study are shown in reagent and tools table.

### siRNA-mediated gene knockdown in myotubes

Gene knockdown was carried out following the manufacturer’s protocol using a 6-well culture plate format (Thermo Fischer Scientific, Waltham, USA). Lipofectamine 3000 was used for the transfection step (Thermo Fischer Scientific, Waltham, USA). C2C12 myotubes and primary human myotubes were transfected with 75 pmol siRNA duplex.

### CRISPR/Cas9-mediated gene overexpression

To induce endogenous overexpression, C2C12 myoblasts were transfected with a DUSP22 CRISPR activation plasmid (Santa Cruz, SC-430587-ACT, sequence: TGCAGTTTGCGCACGCGCGC). The myoblasts were infected for 48 h and then treated with puromycin dihydrochloride (2 µg/ml), hygromycin B (200 µg/ml), and blasticidin S HCl (20 µg/ml) for selection. The expression levels were then confirmed using qPCR to verify overexpression.

### Animal studies

Studies were carried out under the guidance provided by the Institute for Laboratory Animal Research Guide for the Care and Use of Laboratory Animals, and were approved by the Gwangju Institute of Science and Technology Animal Care and Use Committee (study approval number GIST-2021-117). Animal studies have been approved by the appropriate ethics committee and have therefore been performed in accordance with the ethical standards laid down in the 1964 Declaration of Helsinki and its later amendments. Animals were provided by Damool Science, Republic of Korea.

### Dex treatment model of skeletal muscle atrophy and DUSP22 knockdown

The Dex model of muscle atrophy was based on the previously published protocol [68]. In brief, 12 week old male C57BL/6J mice were treated as follows: 1) Vehicle (DMSO in PBS pH 7.4 with 5% Tween 80) alone, 2) 15 mg/kg Dex dissolved in vehicle, 3) 15 mg/kg Dex and 5 mg/kg BML-260 (n=5 per group). Mice were treated daily by intraperitoneal (IP) injection for 14 d and then assessed for muscle condition.

To knockdown DUSP22 expression, intramuscular delivery of DUSP22 siRNA (siRNA ID: 287290, Thermo Fisher Scientific, MA, USA) was carried out using the Invivofectamine 3.0 reagent ((Thermo Fisher Scientific) following the manufacturer’s instructions and previously published protocols [69, 70]. In brief, for the Dex model of muscle wasting, 35 ug siRNA complexes were injected into the TA muscle of 12 weeks-old C57BL/6 mice (n=4 per group), 24 h before IP treatment with 15 mg/kg Dex. Mice were treated with 15 mg/kg Dex every 24 h for 7 days, and received a second injection of 35 ug siRNA complex into the TA muscle at day 3 of the Dex treatment. Muscles were harvested for analysis after 7 days of Dex treatment.

### Skeletal muscle aging model and DUSP22 targeting

To knockdown DUSP22 expression in aging skeletal muscle, 27 months-old male C57BL/6 mice received intramuscular delivery of DUSP22 siRNA using the Invivofectamine 3.0 reagent (n=4 per group). 35 ug siRNA complexes were injected into the TA muscle, followed by a second injection 4 days later. TA muscles were harvested for analysis at day 10 of the experiment.

To assess DUSP22 pharmacological targeting, BML-260 was treated to 24-26 months-old male C57BL/6 mice as follows: (1) vehicle alone (5% DMSO+5% tween 80) and (2) 5 mg/kg BML-260 (n=5 per group). Mice were treated via IP delivery every 24 h for 4 weeks and then assessed for skeletal muscle function and condition.

### Immobilization model

To immobilize the hind limbs, 14-week-old male C57BL/6 mice were anesthetized with isoflurane and both hind limbs were fixed with plastic EP-tubes (Axygen, MCT-175-C). To prevent the tubes from slipping, the junction between the tubes and the hind limbs were wrapped with insulating tape. BML-260 was treated as follows: (1) vehicle alone (5% DMSO+5% tween 80) and (2) 5 mg/kg BML-260 (n=3 per group). The mice were monitored daily and sacrificed after 14 days.

### Grip strength test

Grip strength was measured using the BIO-GS3 device (Bioseb, FL, USA). Mice were placed onto the grid with all four paws attached and gently pulled backwards to measure the grip strength until the grid was released. The maximum grip value, used to represent muscle force, was calculated using 3 trials with an interval of 30 sec.

### Muscle fatigue test

Muscle fatigue was measured the previously described protocol [71], with two different models using the Rotarod. One was the constant model and the other was the accelerating model that is inherent in the rotarod device. In brief, the mice were accommodated to training before the commencement of the fatigue task using an accelerating rotarod (Ugo Basile, Italy). The mice were trained at a constant speed of 13 rpm for 15 min. After a recovery period of 15 min, the mice were placed on the rotarod adjusted to accelerate from 13 to 25 rpm in 3 min for 15 min. 24 h later, the muscle fatigue test was carried out with speeds ramping from 13 to 25 rpm in 3 min, and maintained at 25 rpm for a 30 min interval. Latency to fall off the rotarod was measured for each mouse. A fatigued mouse was classified as falling off 4 times within a 1 min period, which also terminated the test.

### Skeletal muscle dissection and histology

Mice were anesthetized using ketamine (22 mg/kg; Yuhan, Republic of Korea) and xylazine (10 mg/kg; Bayer, Republic of Korea) in saline by IP injection. The quadriceps, gastrocnemius, soleus and TA muscles were then dissected and weighed. For histological analysis, the muscles were fixed by overnight incubation with 4% paraformaldehyde in PBS pH 7.4 at 4 °C. Paraffin sectioning and H&E staining were provided by the Animal Housing Facility, Gwangju Institute of Science and Technology, Republic of Korea (5 µm muscle sections using a Thermo Scientific HistoStar (Thermofisher, USA). H&E staining was carried out with a kit (Merck, Germany)). For immunohistochemistry, TA muscle sections were permeabilized for 15 min (PBS with 0.5% Triton X-100), blocked for 1 h (5% BSA) and incubated with the primary antibody overnight at 4°C. After washing with PBS 3 times for 5 min, the sections were incubated with the secondary antibody at RT for 1 h. The nuclei were counterstained with DAPI in ProLong™ Gold Antifade Mountant (Invitrogen, P36935). Digital images were acquired with Leica DM 2500 (Leica Microsystems). Cross sectional area (CSA) and fiber size distribution was measured using the NIH imaging software Image J 64. To CSA and size distribution of different fiber types was calculated in 4 muscle section as follows: type 2A: 30–40 fibers measured; type 2B: at least 100 fibers measured; type 2X: 30– 40 fibers measured.

### RNA-Seq

RNA samples were harvested from the TA muscles of 24-26 months-old male C57BL/6 mice treated as follows: (1) vehicle alone (5% DMSO+5% tween 80) and (2) 5 mg/kg BML-260. Mice were treated by IP delivery every 24 h for 4 weeks (n=3 per group). RNA-Seq was provided by Macrogen, Republic of Korea. Prior to the commencement of sequencing, QC was carried out using FastQC v0.11.7 (http://www.bioinformatics.babraham.ac.uk/projects/fastqc/). Illumina paired ends or single ends in the sequenced samples were trimmed using Trimmomatic 0.38 with various parameters (http://www.usadellab.org/cms/?page=trimmomatic). The sequences of the samples were mapped and analyzed using HISAT2 version 2.1,0, Bowtie2 2.3.4.1 (https://ccb.jhu.edu/software/hisat2/index.shtml). Potential transcripts and multiple splice variants were assembled using StringTie version 2.1.3b (https://ccb.jhu.edu/software/stringtie/). The normalized sample counts were generated using DESeq2, which was also used for normalization, visualization, and differential analysis. PCA was performed using normalized sample counts. The “apeglm” type was used to calculate shrink log2 fold changes.

### Statistical analysis

The Student’s *t* test or ANOVA with the Dunnett’s test was used to determine statistical significance for two groups, or three or more groups, respectively (GraphPad 7.0 Software, Inc., CA, USA). *p* values of less than 0.05 were deemed to be statistically significant. Unless otherwise stated, experiments were carried out in triplicate and the error bars are presented as mean ± standard error of the mean (SEM[*in vivo*]), standard deviation(SD[*in vitro]*)

### Data availability

RNA-Seq data: Gene Expression Omnibus GSE111016 (https://www.ncbi.nlm.nih.gov/geo/query/acc.cgi?acc=GSE111016) RNA-Seq data: Gene Expression Omnibus GSE117525 (https://www.ncbi.nlm.nih.gov/geo/query/acc.cgi?acc=GSE117525)

## Author contributions

S-H. L. carried out experiments, analyzed the data and wrote the manuscript. S-H. L., H-J. K. and S-W. K. carried out experiments. H. L. analyzed the data. D-W. J. and D. R. W. acquired funding, supervised the research, and wrote the manuscript. Conflict of interest: S-H. L, H. L., D-W. J. and D. R. W. are named co-inventors of a pending provisional patent application based in part on the research reported in this paper.

---

**Supplementary Table 1.** Primary antibodies used in this study.

| Primary antibody | Clone | Company | Catalog No. | Dilution |
| --- | --- | --- | --- | --- |
| **α-Tubulin** | Polyclonal | INVITROGEN | PA5-29444 | 1:10000 |
| **Atrogin-1/MAFbx** | Monoclonal | Abcam | Ab168372 | 1:1000 |
| **MuRF-1/Trim63** | Monoclonal | SANTA CRUZ | SC-398608 | 1:500 |
| **FoxO3a** | Monoclonal | Cell Signaling | #12829 | 1:1000 |
| **Phospho-FoxO3a** | Polyclonal | Cell Signaling | #9466 | 1:1000 |
| **AKT** | Monoclonal | Cell Signaling | #2983 | 1:1000 |
| **Phospho-AKT** | Monoclonal | Cell Signaling | #2971 | 1:1000 |
| **JNK** | Monoclonal | Cell Signaling | SC-7345 | 1:1000 |
| **Phospho-JNK** | Monoclonal | Cell Signaling | #4668 | 1:1000 |
| **c-Jun** | Monoclonal | Cell Signaling | #9165 | 1:1000 |
| **Phospho-c-Jun** | Monoclonal | Cell Signaling | #2361 | 1:1000 |
| **STAT3** | Monoclonal | SANTA CRUZ | SC-8019 | 1:1000 |
| **Phospho-STAT3** | Monoclonal | SANTA CRUZ | SC-8059 | 1:1000 |
| **Myosin heavy chain2** | Monoclonal | SANTA CRUZ | SC-53095 | 1:1000 |
| **DUSP22** | Polyclonal | Abcam | Ab70124 | 1:1000 |
| **Myosin Heavy Chain 2 A** | Monoclonal | DSHB | SC-71 | 2ug/ml |
| **Myosin Heavy Chain 2 B** | Monoclonal | DSHB | BF-F3 | 2ug/ml |
| **Laminin** | Polyclonal | sigma | L9393 | 1:50 |
| **Fast-type skeletal muscle** | monoclonal | SANTA CRUZ | sc-32732 | 1:100 |

**Supplementary Table 2.** Secondary antibodies used in this study.

| Primary antibody | Conjugated use | Company | Catalog No. | Dilution |
| --- | --- | --- | --- | --- |
| Goat anti-Mouse IgG (H+L) | HRP | Abcam | ab6789 | 1:10000 |
| Goat anti-Rabbit IgG HRP | HRP | Cell signaling | #7074S | 1:10000 |
| Alexa Fluor™ 488 Goat anti-mouse IgG(H+L) | Alexa Fluor 488 | INVITROGEN | A11001 | ICC:1:2000 |
| Alexa Fluor™ 555 Goat anti-mouse IgG(H+L) | Alexa Fluor 555 | INVITROGEN | A21422 | IHC 1:500 |
| Alexa Fluor™ 488 Goat anti-mouse IgG(H+L) | Alexa Fluor 488 | INVITROGEN | A28175 | IHC 1:500 |
| Alexa Fluor™ 350 Goat anti-mouse IgG(H+L) | Alexa Fluor 350 | INVITROGEN | A11045 | IHC 1:500 |

**Supplementary Table 3.**
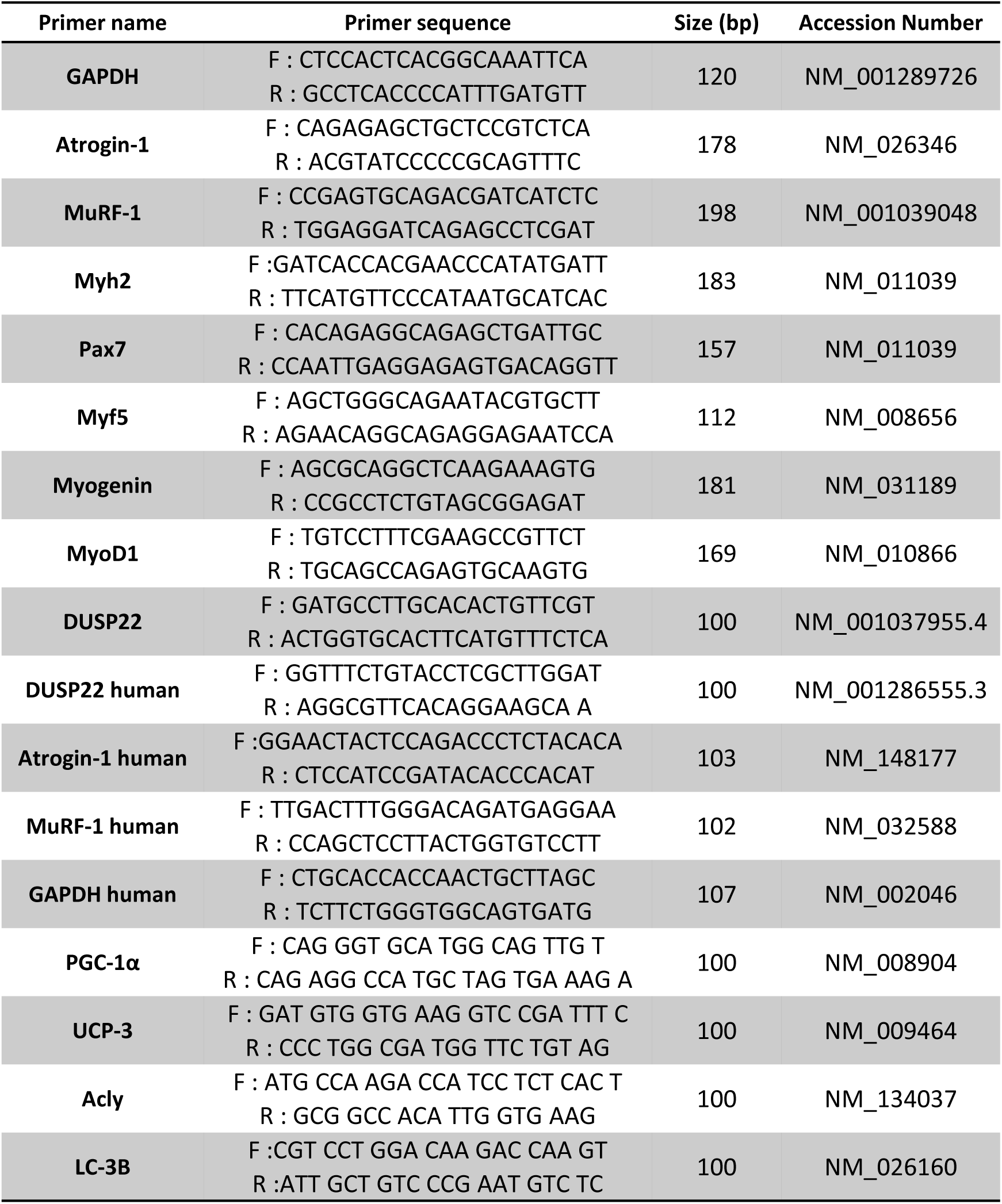
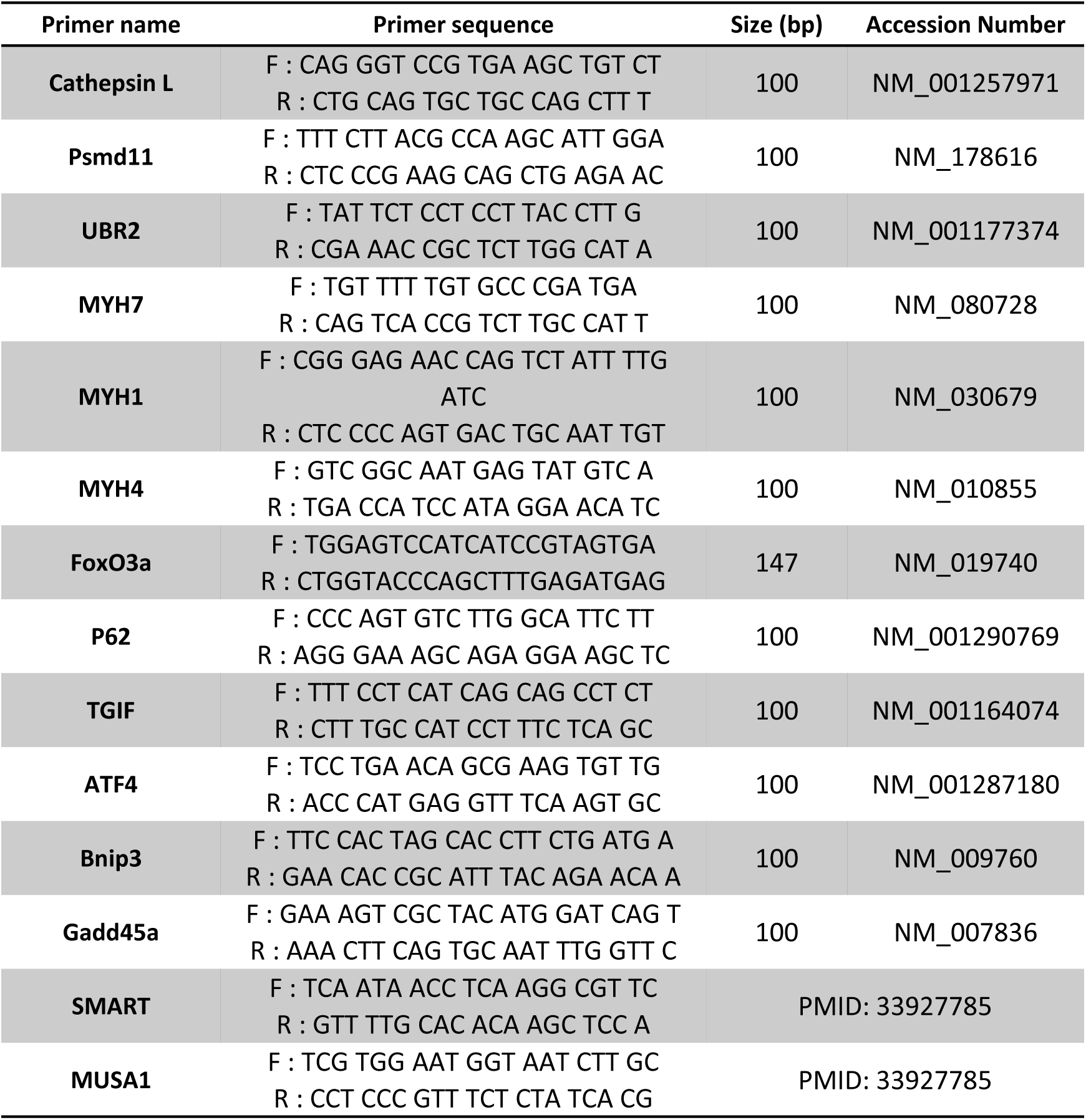
Primers used for qPCR.

